# Sugar alcohols have the potential as bee-safe feeding stimulants for wasp control

**DOI:** 10.1101/2022.02.04.479032

**Authors:** Stefanie Neupert, Jennifer M. Jandt, Paul Szyszka

## Abstract

Pest insects are often baited with poisoned feeding stimulants, the most common of which are sugars. However, sugars are attractive for most animal species, which makes it difficult to target only a specific pest insect species. Here, we assessed different sugar alcohols for their potential as more species-selective feeding stimulants for pest insects. We tested the attractiveness of the sugar alcohols sorbitol, xylitol and erythritol with a capillary feeder assay in wasps (as potential pest insects, because introduced wasps are a pest in many regions) and bees (as non-target insects). For the common wasp (*Vespula vulgaris*), sorbitol and xylitol acted as nutritive feeding stimulants, and erythritol acted as a non-nutritive feeding stimulant. For the buff-tailed bumble bee (*Bombus terrestris*), sorbitol acted as a feeding stimulant, while for the honey bee (*Apis mellifera*), none of the sugar alcohols acted as feeding stimulant. The species-specific preferences for sugar alcohols suggest their potential as species-selective insect baits. The wasp-specific preference for xylitol suggests its potential as bee-safe alternative to sugar-containing bait for wasp pest control.

**GRAPHICAL ABSTRACT:** We tested the attractiveness of sugar alcohols with a capillary feeder assay in wasps and bees. Species-specific preferences suggests that sugar alcohols have the potential for being used as bee-safe insect baits.

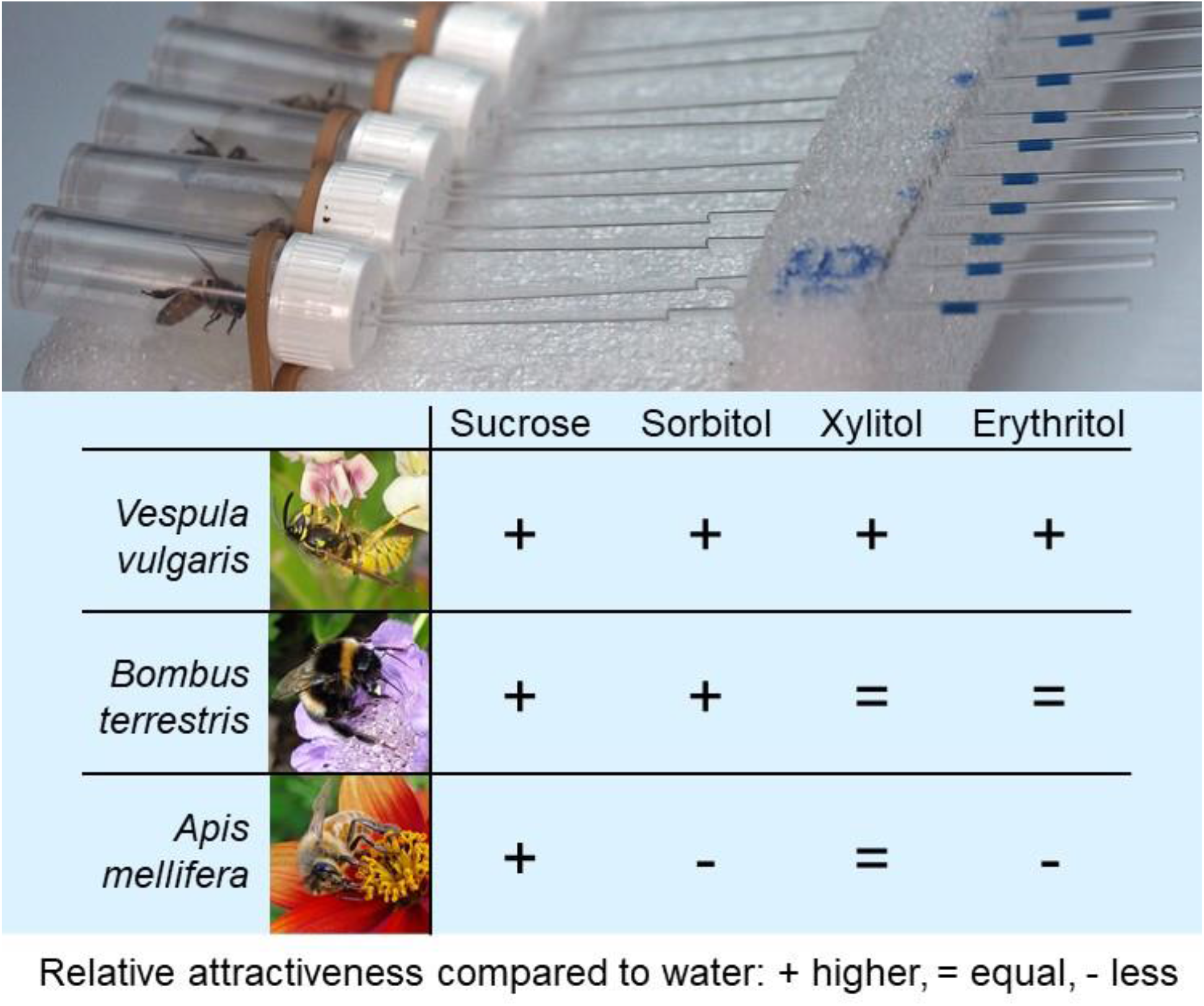

## 1 INTRODUCTION

An efficient and sustainable pest control should target a given pest species while minimizing impact on non-target species. Pest insects are often baited with poisoned feeding stimulants. The most common feeding stimulants are sugars (e.g., sucrose, glucose, fructose)^1^. However, it is difficult to hold off non-target animals from sugar baits because sugar preference is not species-selective (most insect species consume sugars)^2^. Fats and proteins are more species-selective feeding stimulants compared to sugars^1^. Sugar-free protein baits, for example, are readily consumed by social wasps^3,4^, but not by honey bees^5^. A drawback of using protein baits on pest insects, however, is that many insects prefer proteins over sugars only during the limited period of reproduction^4,6^. In contrast to proteins, most insects consume carbohydrates (e.g., sugars) all year round to meet their daily energy requirements^2^.

The aim of this study was to assess non-sugar carbohydrates for their potential as a species-selective bait for insects. We tested honey bees (*Apis mellifera*) and buff-tailed bumble bees (*Bombus terrestris*) as representatives of non-target insects. We tested common wasps (*Vespula vulgaris*) as a representative of a potential pest insect, because *Vespula* social wasps are considered a nuisance in most parts of the world^7^, and are a major ecological and economic pest in regions they have invaded^8^. As non-sugar carbohydrates we used the odourless sugar alcohols sorbitol, xylitol and erythritol, which occur naturally in fruits (sorbitol^9,10^ and xylitol^10^), honeydew (sorbitol^11^) and fermented fruits (erythritol^12^), and are used as human-safe sweeteners^13^. We chose these sugar alcohols, because they are feeding stimulants for some, but not all, insect species^14^. Sorbitol and erythritol are potential candidates for bee-safe insect baits, because honey bees only consume them when mixed with an appetitive substance such as sucrose^15^. To our knowledge, there are no studies on xylitol preferences in honey bees, or on sorbitol, xylitol or erythritol preferences in bumble bees or *Vespula* wasps.

We tested the attractiveness of sorbitol, xylitol and erythritol with a two-choice capillary feeder assay similar to the feeder used in May et al.^16^. We found that for wasps, sorbitol and xylitol acted as nutritive feeding stimulants and erythritol acted as a non-nutritive feeding stimulant. For bumble bees, sorbitol acted as a feeding stimulant, and for honey bees, none of the sugar alcohols acted as feeding stimulant. We propose xylitol as a candidate for bee-safe carbohydrate baits for the common wasp and other pest insects.

## 3 MATERIALS AND METHODS

### 3.1 Animals

We used female workers of the common wasp (*Vespula vulgaris*; 326 individuals), the buff-tailed bumble bee (*Bombus terrestris*; 149 individuals), and the honey bee (*Apis mellifera*; 110 individuals). Wasps were caught in March and April 2021 from their nest entrance from four colonies in and around Dunedin, New Zealand. Wasps were collected into 300 ml plastic containers, transported to the lab on the same day and kept in the fridge at 5°C without food for up to 3 days (wasps can be kept alive in the fridge without food for at least 4 days (JM Jandt, unpublished data)). Bumble bees were collected from six colonies that were purchased from Biobees Ltd, Hastings, New Zealand. Before collection, bumble bee colonies were kept in wooden nest boxes in a greenhouse at Invermay Research Centre, Mosgiel, where they were allowed to forage on tomato (pollen only), borage (pollen and nectar) and 1.5M sucrose solution. Honey bees were caught from one bee hive on the roof top of the Zoology Department, University of Otago.

### 3.2 Two-choice capillary feeder assay

Animals were cooled on ice until they stopped moving, and each individual was put in a transparent cylindric plastic chamber (14 mm inner diameter, 50 mm length; screw cap vials, Thermo Fisher) and placed in a horizontal position. The bottom end had a 3 mm hole for oxygen exchange. The screw cap had two 3 mm holes, side by side and 6 mm apart from each other for inserting calibrated glass capillaries (100 μl micropipettes, 1.7 mm outer diameter; VWR). Capillaries were filled with 100 μl tastant solution or water. The sides (left or right) of the capillaries containing the tastant solution or water were balanced across animals to cancel out potential side-preferences. Approximately 1 hour after animals were put into their chambers, unresponsive or sluggish animals were discarded. Capillaries were inserted 5 mm into the chamber and angled 5° relative to the horizontal, so that the liquid would flow towards the animal without leaking out of the capillary. Up to 10 chambers (i.e., 10 animals) were placed into a 3.5 litre transparent plastic box equipped with five 10 mm holes for air exchange, and with a water-soaked tissue to get approximately 85% relative humidity. Those boxes were placed in a laboratory at 22°C, with natural light through the window, but not exposed to direct sunlight. Experiments started between 15:00 and 19:00 hours. After the animal had been in the chamber for 24 hours, we recorded if the animal was alive or dead, and we measured the volume of liquid that remained in the capillaries with a calliper (75 mm capillary length corresponded to 100 μl and the measuring accuracy was ± 1 mm 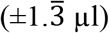).

To test the animals’ preference for sucrose and sugar alcohol solutions over water (Fig. 1, data set 1), we provided each animal with a choice of either sucrose or sugar alcohol solution as tastant solution and distilled water as control. To test, if wasps consume sorbitol or xylitol solution in presence of sucrose solution (Fig. 2, data set 2), we provided them with a choice of either sorbitol or xylitol as tastant solution and one of four concentrations of sucrose solutions as control. To measure wasps’ survival rate on water, and test for potential side preferences, we provided them with water in both capillaries (Fig. S1, data set 3).

**Figure 1.**
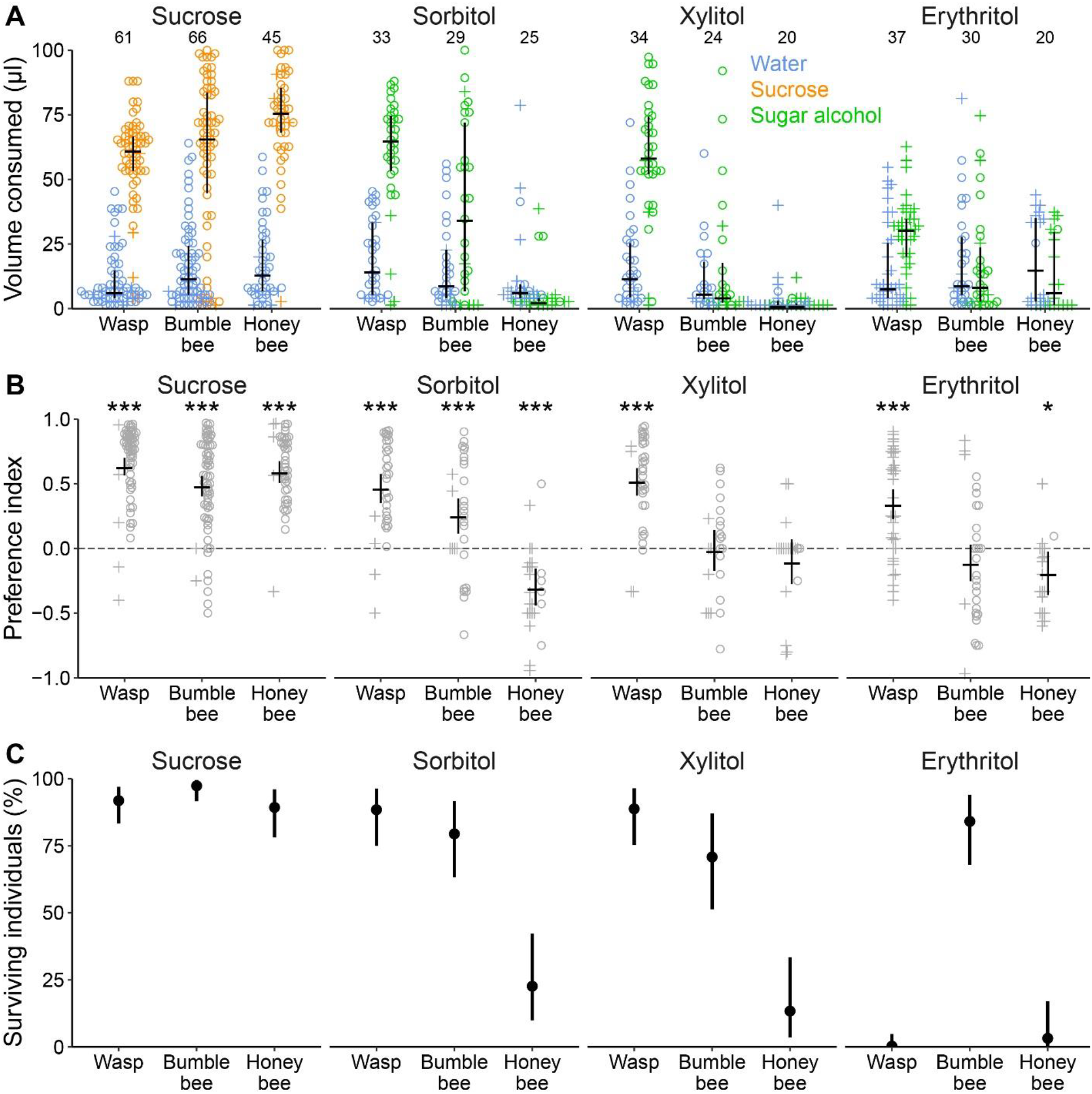
Sugar alcohols are feeding stimulants for the common wasp but not for the honey bee. **A**) Sucrose (1.25M), sorbitol (1.7M), xylitol (1.5M) and erythritol (1.5M) solution and water consumption in a two-choice capillary feeder assay. Volume of sucrose/sugar alcohol solution and water consumed during 24h. Colors indicate solution type (blue = water, orange = sucrose, green = sugar alcohol). Symbols indicate if an individual was alive (circles) or dead (crosses) after 24h. Horizontal lines indicate the median, and vertical lines indicate 25% and 75% quantiles. Numbers above data points indicate the number of animals tested in that treatment group (each animal is represented by two data points, one for sucrose/sugar alcohol solution and one for water). **B**) Preference index for the data in A. Positive preference indices represent a preference for sucrose or sugar alcohol solution, and negative preference indices represent a preference for water. Horizontal black lines indicate estimated averages of preference indices, and vertical black lines indicate 95% credible intervals. Asterisks indicate the certainty that the preference index is different to zero (i.e., there is a preference for either the tastant solution or water): * >95%, *** >99.9% certainty. **C**) Survival after 24h differs between species and depends on the type of solution that was offered as alternative to water. Dots indicate estimated averages of survival rate, and vertical black lines indicate 95% credible intervals.

**Figure 2.**
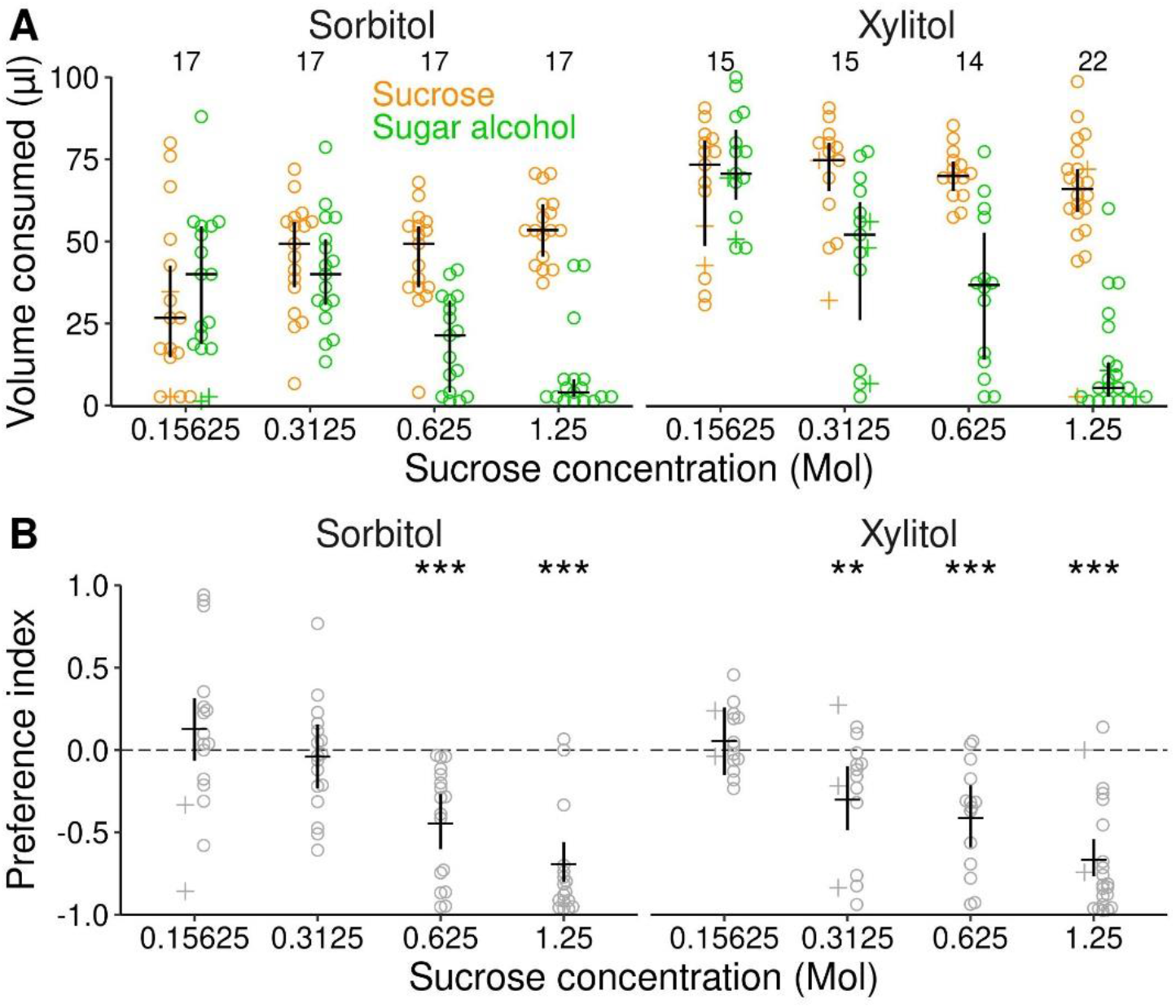
Wasps consume sorbitol and xylitol when sucrose is available. **A**) Sugar alcohol solution and sucrose consumption in a two-choice capillary feeder assay. Volume of sucrose (concentration varied) and sorbitol solution (1.7M) or xylitol solution (1.5M) consumed during 24h. Colors indicate solution type (orange = sucrose, green = sugar alcohol). Numbers above data points indicate number of animals tested in that treatment group (each animal is represented by two data points, one for sucrose solution and one for sugar alcohol solution). The volume of consumed sugar alcohol solution decreased with increasing sucrose concentration. **B**) Preference index for the data in A. Negative preference indices represent a preference for sucrose over the sugar alcohol. Asterisks indicate the certainty that the preference index is different to zero (i.e., there is a preference for the sucrose solution over the sugar alcohol solution): ** >99%, *** >99.9% certainty.

### 3.3 Tastant solutions

We created the tastant solutions by dissolving tastants in distilled water at the following concentrations: Sucrose, 1.25M (Chelsea Sugar, New Zealand), erythritol, 1.5M (Active Bio-Tech Limited, New Zealand), sorbitol 1.7M (Sigma Aldrich, USA), and xylitol 1.5M (Xlear, New Zealand). The concentrations of sucrose and sorbitol were chosen to achieve similar nutritive values for honey bees^17^. The concentration of erythritol and xylitol were similar as in previous studies in different insect species^14^. For data set 2, we created three serial dilutions of the 1.25M sucrose solution to achieve 0.625M, 0.3125M and 0.15625M solutions.

### 3.4 Data Analysis

#### 3.4.1 Volumes consumed

First, we calculated the volume of the two liquids consumed by each animal tested. Therefore, we calculated the difference between the liquid column in the capillaries at the beginning of the experiment (75 mm) and after 24h. The difference in the liquid column was multiplied by 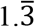 to transform the difference of liquid column in mm to the corresponding volume consumed in μl. To avoid division through zero when calculating the preference index (3.4.2), consumed volumes of zero were set to 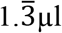 (1 mm corresponds to 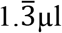). We used those adjusted volumes consumed in plots and for further analysis (hereafter referred to as consumed volumes). To visualize the consumed volumes of the provided liquids, we plotted the single data points with the median and 50% quantiles of each of the two provided liquids consumed for each experimental group (Figures 1A and 2A).

#### 3.4.2 Preference index

As a measure for the individual preference for the two liquids provided in the two-choice capillary feeder assay, we calculated a preference index^18^:

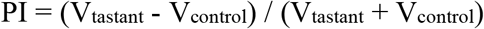

where V_tastant_ is the consumed volume of tastant solution and V_control_ is the consumed volume of control solution. Generally, a preference index can range from −1 to +1, where a preference index of +1 indicates that only tastant solution was consumed while a preference index of −1 indicates that only the control solution was consumed. Note that in our analysis, the preference indices can only range from −0.97 to +0.97 because consumed volumes of zero were set to 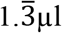 (see 3.4.1). A preference index of zero indicates that equal volumes of tastant and control solution were consumed.

To estimate the average preference index for each experimental group, we rescaled the preference index to range from 0 to 1.

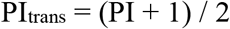

where PItrans is the rescaled (transformed) preference index.

#### 3.4.3 Statistics

To estimate the species-specific preference indices for the different tastants (data set 1; Fig. 1B), we ran a beta regression model using the *stan_betareg* function with the transformed preference index (ranging between 0 and 1) as response variable, and *species* and *tastant* as explanatory variables including an interaction term. To estimate the survival probability in data set 1 (Fig. 1C), we ran a binomial model using the *stan_glm* function with survival *yes* or *no* (1/0) as the binary response variable and *species* and *tastant* as explanatory variables including an interaction term. To estimate the preference indices of wasps for different sucrose concentrations (data set 2; Fig. 2B), we ran a beta regression model with the transformed preference index as the response variable. We added *tastant* and *sucrose concentration* as explanatory variables and included an interaction term. To test if wasps had a preference for one side over the other (data set 3, Fig. S1), we ran a beta regression model using the *stan_betareg* function and added only the preference index as response variable. To estimate the survival probability in data set 3 (Fig. S1C), we ran a binomial model using the *stan_glm* function with survival *yes* or *no* (1/0) as the binary response variable only.

All models were run in a Bayesian framework using *rstanarm* R package utilities^19^. We used the default weakly informative priors of a normal distribution with a mean of 0 and a scale of 2.5. We set the number of chains to 5, each chain with 20,000 iterations. The first 10,000 interations were used as burn-in. For all models, we did graphical posterior predictive model checking. In addition, we checked that all Rhat values were close to one showing no indication for non-convergence. We also checked that effective sample sizes were higher than 10,000 to yield stable estimates for the 95% credible intervals. To draw inferences, we drew 20,000 random samples from the posterior distribution of the model parameters and used the 2.5% and 97.5% quantiles as the lower and upper limits of the 95% credible intervals. To calculate the certainty of a preference index being different to zero, for each experimental group we assessed the proportion of random values being higher than 0.5 (note that 0.5 corresponds to a preference index of zero in the original scale). If the proportion of values higher than 0.5 was above 0.95 (or below 0.05), we can be 95% certain that the PI is higher than 0 (or lower than 0, respectively). For plotting, we back-transformed the preference indices and the credible intervals into the original scale (−1, +1).

All data were analyzed in R v4.1.0^20^ and all plots were created with the *ggplot2* package.

## 2 RESULTS

To validate the suitability of the feeder assay for measuring the attractiveness of sugar alcohols, we measured the attractiveness of sucrose solution (1.25M) relative to water, because sucrose is a strong feeding stimulant for the common wasp^21^, the buff-tailed bumble bee^22^, and the honey bee^23^. Across all species, most individuals consumed sucrose solution (averages between 61 to 76 μl) but little or no water (1 to 15 μl) (Fig. 1A). All species preferred sucrose over water (average preference indices between 0.49 and 0.64) (Fig. 1B), confirming the suitability of the feeder assay for measuring the attractiveness of tastants.

### 2.1 Wasps and bees differ in their preferences for sugar alcohols

Sorbitol solution (1.7M) acted as a feeding stimulant for wasps and bumble bees (Fig. 1A), which preferred sorbitol over water (Fig. 1B). However, sorbitol preference was higher in wasps (average preference index 0.47) than in bumble bees (0.25). In contrast, most honey bees consumed little, or no sorbitol (Fig. 1A), and preferred water over sorbitol (average preference index −0.3, Fig. 1B).

Xylitol solution (1.5M) acted as feeding stimulant for wasps (Fig. 1A) and they preferred it over water (Fig. 1B). Most bumble bees and honey bees consumed little, or no xylitol, and they did not differentiate between xylitol and water (average preference index not different from zero).

Erythritol solution (1.5M) acted as feeding stimulant for wasps (Fig. 1A), and they preferred it over water (Fig. 1B), but their preference for erythritol was lower (0.34) than their preference for sucrose (0.64). Bumble bees did not differentiate between erythritol and water, and honey bees preferred water over erythritol.

None of the erythritol-fed wasps survived (Fig. 1C), and only 11% of water only-fed wasps survived (three out of 27 wasps, Fig. S1). This indicates that erythritol is non-nutritive. In contrast, sorbitol- and xylitol-fed wasps survived equally well as the sucrose-fed wasps. This indicates that sorbitol and xylitol are attractive and nutritive for wasps.

### 2.2 Wasps consume sorbitol and xylitol in the presence of sucrose solution

Because in the wild, wasps forage on sucrose-containing food sources (e.g., fruits, nectar, honeydew), we next tested whether wasps consume sorbitol and xylitol when sucrose is available. Wasps preferred sucrose over sorbitol when the sucrose concentration was high (0.625M or 1.25M (Fig. 2)). However, when the sucrose concentration was reduced to 0.3125M or 0.15625M, wasps consumed equal volumes of sorbitol and sucrose solution (Fig. 2B). Likewise, wasps preferred sucrose over xylitol when the sucrose concentration was 0.3125M or higher (Fig. 2). At the lowest sucrose concentration (0.15625M), wasps consumed equal volumes of xylitol and sucrose solution (Fig. 2B).

## 4 DISCUSSION

Sugars are the most frequently used carbohydrates in baits for controlling pest insects^1^. However, sugar baits are also attractive to many non-target insects^2^. Here, we assessed non-sugar carbohydrates, sugar alcohols, for their potential to serve as bee-safe feeding stimulants for pest insect control. We found that sorbitol, xylitol and erythritol acted as feeding stimulants for wasps, but not for honey bees, and sorbitol acted as a feeding stimulant for bumble bees, but its action was weak.

### 4.1 Sugar alcohols differ in their attractiveness, nutritive value and toxicity between insect species

Sorbitol is used in pest insect baits where it serves as an humectant to maintain the bait’s moisture (e.g., in sugar-baits for cockroaches^24^ or in protein-baits for *Vespula* wasps^25^). Sorbitol acts as a feeding stimulant for some insect species (fruit flies^26,27^, some ant ^28^ and cockroach species^29^), but not for others (honey bee^17^). Our data confirms the species-specificity of sorbitol attraction: Sorbitol acted as feeding stimulant for common wasps, to a lesser extent for bumble bees, but not for honey bees, who even preferred water over sorbitol solution. This fact, that for all tested insect species sorbitol is nutritive and non-toxic (honey bee^17^, some fly species^27,30–37^, yellow fever mosquito^32^, German cockroach^38^, a solitary wasp^39^), corresponds to the high survival rate of sorbitol-fed wasps in our study. Remarkably, honey bees starved to death rather than consuming sorbitol which would likely have saved their lives, given that sorbitol is nutritive for honey bees^17^ and that in our study honey bees survived on a sucrose diet.

Xylitol and erythritol are toxic to several insect species and their use as insecticides has been suggested^36,40,49,50,41–48^. Xylitol consumption has only been tested in flies. Xylitol is a feeding stimulant, nutritive but toxic for the house fly^42^, nutritive for some *Drosophila* species^35,36^, and toxic for the stable fly^43^. We found that xylitol acted as a feeding stimulant for the common wasp, but not for bumble bees or honey bees. The high survival rate of xylitol-fed wasps suggests that xylitol is nutritive.

Erythritol is a feeding stimulant for the fruit fly^40^ and for the house fly^42^, but it is not a feeding stimulant for the spotted wing *Drosophila^36^*, for several ant species^28^ or for the honey bee^17^. Likewise, erythritol preferences differed across wasps and bees: Erythritol acted as a feeding stimulant in common wasps but not in bumble bees, and honey bees preferred water over erythritol solution. Erythritol is non-nutritive for all tested insect species (the honey bee^17^ and several fly species^30,31,34,36,37^), and it is toxic for several insect species (some fly species ^36,40–45^, yellow fever mosquito^46^, some ant species^47,48^, pear psylla^49^, eastern subterranean termite^50^). Correspondingly, the low survival rate of erythritol-fed wasps suggests that erythritol is non-nutritive for the common wasp. A previous study found that erythritol is not toxic for honey bees^17^. Therefore, the low survival rate of erythritol-fed honey bees likely reflects starvation due to the lack of consumption or lack of nutritive value rather than a toxicity of erythritol.

### 4.2 Sorbitol, xylitol and erythritol have the potential for being used as bee-safe wasp baits

*Vespula* social wasps are an ecological and economic pest in large parts of the southern hemisphere^8^. Vespula wasps are baited with sugar-free protein baits, which they readily consume in spring and early summer when the colony is collecting protein for brood production^3,4,6^. Bumble bees and honey bees also rely on protein for brood development, but they receive proteins from pollen and are not attracted to the protein baits developed for wasps^5^. A drawback of using protein baits on wasps, however, is that the window of opportunity to use these baits is limited to early in the colony development, when colony sizes are still small and less of a nuisance. By late summer or early autumn, colony sizes reach their peaks and workers shift their foraging preference to carbohydrates (e.g., sugars)^6^. At this time, a protein bait becomes ineffective.

Our data show that the carbohydrates sorbitol, xylitol and erythritol are feeding stimulants for the common wasp but not for the honey bee. Bumble bees showed a preference for sorbitol over water, but not for the other sugar alcohols. Therefore, sorbitol may be used as bee-safe bait in areas where there are honey bees but no bumble bees. Xylitol appears to be the most promising candidate for a bee-safe wasp bait, because it is neither attractive for the buff-tailed bumble bee nor for the honey bee, and because it has nutritive value for the common wasp.

What are the next steps in assessing these sugar alcohols for their use as pest insect baits? We found that wasps consume 1.5M sorbitol or xylitol solution in the presence of 0.625M sucrose solution, but not in the presence of 1.25M sucrose solution. Natural sugar sources contain sugar concentrations that can be equivalent to or higher than 1.25M of sucrose (e.g., nectar^51^, honeydew^52^, fruits^53^). Therefore, future research should investigate at what concentrations and in which environments sugar alcohols can effectively attract wasps (and potentially other pest insects), considering the natural sugars that are available at those places and times.

## ACKNOWLEDGEMENTS

We thank Melita Busch and Mateus Detoni for collecting wasps, and Mateus Detoni, Jack Matterson and Melita Busch for giving feedback to this manuscript.

## DATA AVAILABILITY

Data and analysis code will be made available on Open Science Framework (https://osf.io/)

## SUPPORTING INFORMATION

**Figure S1.**
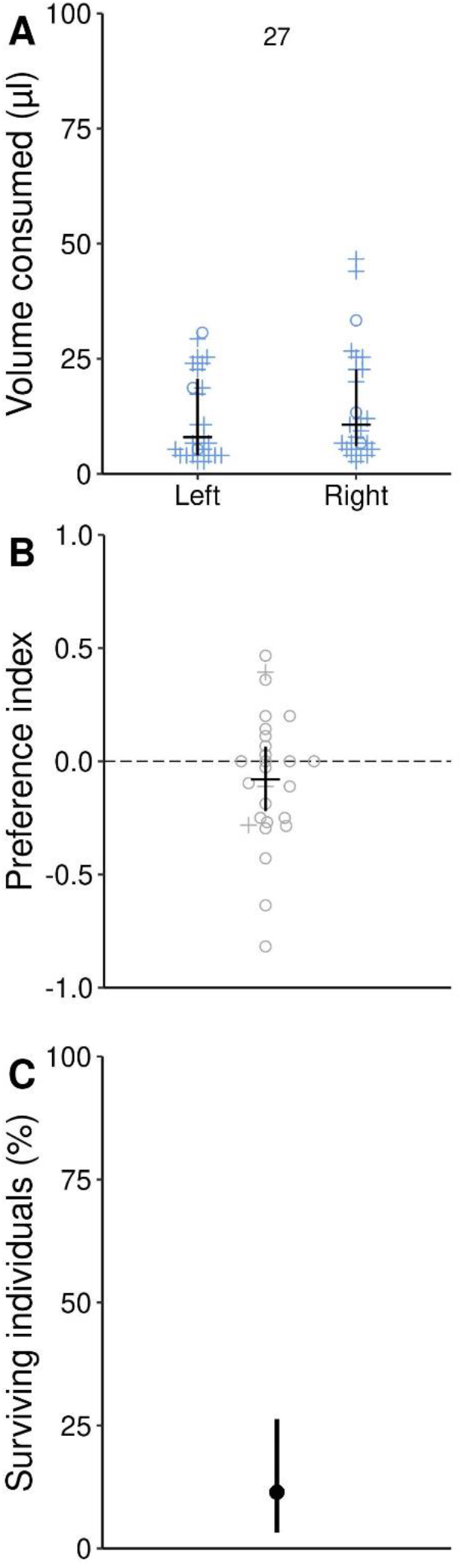
Water only-fed wasps do not prefer one side over the other and only few individuals survive. **A**) Water consumption in a two-choice capillary feeder assay where both, the left and right capillary contained water. Symbols indicate if an individual was alive (circles) or dead (crosses) after 24h. Horizontal lines indicate the median, and vertical lines indicate 25% and 75% quantiles. Number above datapoints indicate number of animals (each animal is represented by two data points, one for each capillary). **B**) Preference index for the data in A. Positive preference indices represent a preference for water on the left side, and negative preference indices represent a preference for water on the right side. The horizontal black line indicates the estimated average of the preference index and the vertical line indicates 95% credible interval. The preference index was −0.08 (credible interval: −0.22 to 0.07) indicating that wasps had no preference for one of the sides. **C**) Survival after 24h was low (three out of 27 individuals survived). Dot indicates estimated average of preference index and vertical line indicates 95% credible interval.

## REFERENCES

1. Avé, D. A. Stimulation of Feeding: Insect Control Agents. Regul. Mech. Insect Feed. 345–363 (1995). doi:10.1007/978-1-4615-1775-7_12

2. Stoffolano, J. G. Regulation of a Carbohydrate Meal in the Adult Diptera, Lepidoptera, and Hymenoptera. in Regulatory Mechanisms in Insect Feeding 210–247 (Springer US, 1995). doi:10.1007/978-1-4615-1775-7_8

3. Edwards, E., Toft, R., Joice, N. & Westbrooke, I. The efficacy of Vespex® wasp bait to control Vespula species (Hymenoptera: Vespidae) in New Zealand. https://doi.org/10.1080/09670874.2017.1308581 63, 266–272 (2017).

4. Richter, M. R. Social Wasp (Hymenoptera: Vespidae) Foraging Behavior. http://dx.doi.org/10.1146/annurev.ento.45.1.121 45, 121–150 (2003).

5. Edwards, E. D., Woolly, E. F., McLellan, R. M. & Keyzers, R. A. Non-detection of honeybee hive contamination following Vespula wasp baiting with protein containing fipronil. PLoS One 13, (2018).

6. Browne, L. B. Physiologically induced changes in resource-oriented behavior. Annu. Rev. Entomol. Vol 38 1–25 (1993). doi:10.1146/annurev.ento.38.1.1

7. Sumner, S., Law, G. & Cini, A. Why we love bees and hate wasps. Ecol. Entomol. 43, 836–845 (2018).

8. Lester, P. J. & Beggs, J. R. Invasion success and management strategies for social vespula wasps. Annual Review of Entomology 64, 51–71 (2019).

9. Bieleski, R. L. Sugar Alcohols. Plant Carbohydrates I 158–192 (1982). doi:10.1007/978-3-642-68275-9_5

10. Mäkinen, K. K. & Söderling, E. A quantitative study of mannitol, sorbitol, xylitol, and xylose in wild berries and commercial fruits. J. Food Sci. 45, 367–371 (1980).

11. Dhami, M. K., Gardner-Gee, R., van Houtte, J., Villas-Bôas, S. G. & Beggs, J. R. Species-specific chemical signatures in scale insect honeydew. J. Chem. Ecol. 37, 1231–1241 (2011).

12. Shindou, T. et al. Identification of erythritol by HPLC and GC-MS and quantitative measurement in pulps of various fruits. J. Agric. Food Chem. 37, 1474–1476 (2002).

13. Grembecka, M. Sugar alcohols—their role in the modern world of sweeteners: a review. Eur. Food Res. Technol. 241, 1–14 (2015).

14. Lee, S. H., Choe, D. H. & Lee, C. Y. The Impact of Artificial Sweeteners on Insects. J. Econ. Entomol. 114, 1–13 (2021).

15. Frisch, K. v. Versuche über den Geschmackssinn der Bienen. Naturwissenschaften 16, 307–315 (1928).

16. May, P. G. A simple method for measuring nectar extraction rates in butterflies. J. Lepid. Soc. 39, 53–55 (1985).

17. Vogel, B. Über die Beziehungen zwischen Süßgeschmack und Nährwert von Zuckern und Zuckeralkoholen bei der Honigbiene. Zeitschrift für vergleichende Physiol. 1931 142 14, 273–347 (1931).

18. Quinn, W. G., Harris, W. A. & Benzer, S. Conditioned behavior in Drosophila melanogaster. Proc. Natl. Acad. Sci. U. S. A. 71, 708–12 (1974).

19. Goodrich B, Gabry J, Ali I & Brilleman S. rstanarm: Bayesian applied regression modeling via Stan. R package version 2.21.1, https://mc-stan.org/rstanarm (2020). Available at: https://mc-stan.org/rstanarm. (Accessed: 17th January 2022)

20. R Core Team. R Core Team (2021). R: A language and environment for statistical computing. R Foundation for Statistical Computing, Vienna, Austria. URL https://www.R-project.org/ - Google Suche. R Foundation for Statistical Computing, Vienna, Austria. URL https://www.R-project.org/ (2021). Available at: https://www.r-project.org/. (Accessed: 17th January 2022)

21. Free, J. B. The behaviour of wasps (Vespula germanica L. and V. Vulgaris L.) when foraging. Insectes Soc. 17, 11–19 (1970).

22. Mommaerts, V., Wäckers, F. & Smagghe, G. Assessment of Gustatory Responses to Different Sugars in Harnessed and Free-Moving Bumblebee Workers (Bombus terrestris). Chem. Senses 38, 399–407 (2013).

23. v. Frisch, K. Versuche über den Geschmackssinn der Bienen. Naturwissenschaften 15, 321–327 (1927).

24. Schal, C. & Hamilton, R. L. Integrated suppression of synanthropic cockroaches. Annu. Rev. Entomol. 35, 521–551 (1990).

25. Harris, R. J. & Rees, J. S. Aerial poisoning of wasps. Sci. Conserv. 1–26 (2000).

26. Gordesky-Gold, B., Rivers, N., Ahmed, O. M. & Breslin, P. A. S. Drosophila melanogaster prefers compounds perceived sweet by humans. Chem. Senses 33, 301–309 (2008).

27. Fujita, M. & Tanimura, T. Drosophila evaluates and learns the nutritional value of sugars. Curr. Biol. 21, 751–755 (2011).

28. Schmidt, A. Geschmacksphysiologische Untersuchungen an Ameisen. Zeitschrift für vergleichende Physiol. 1938 253 25, 351–378 (1938).

29. Tsuji, H. Studies on the behaviour pattern of feeding of three species of cockroaches, Blattella germanica (L.), Periplaneta americana L., and P. fuliginosa S., with special reference to their responses to some constituents of rice bran and some carbohydrates. Med. Entomol. Zool. 16, 255–262 (1965).

30. Fraenkel, G. Utilization and Digestion of Carbohydrates by the Adult Blowfly. J. Exp. Biol. 17, 18–29 (1940).

31. Hassett, C. C., Dethier, V. G. & Gans, J. A comparison of nutritive values and taste thresholds of carbohydrates for the blowfly. Biol. Bull. 99, 446–453 (1950).

32. Galun, R. & Fraenkel, G. Physiological effects of carbohydrates in the nutrition of a mosquito, Aedes aegypti and two flies, Sarcophaga bullata and Musca domestica. J. Cell. Comp. Physiol. 50, 1–23 (1957).

33. Burke, C. J. & Waddell, S. Remembering nutrient quality of sugar in Drosophila. Curr. Biol. 21, 746–50 (2011).

34. Hassett, C. C. The utilization of sugars and other substances by Drosophila. Biol. Bull. 95, 114–123 (1948).

35. Kircher, H. W. & Al-Azawi, B. Longevity of seven species of cactophilic Drosophila and D. melanogaster on carbohydrates. J. Insect Physiol. 31, 165–169 (1985).

36. Choi, M. Y. et al. Effect of non-nutritive sugars to decrease the survivorship of spotted wing drosophila, Drosophila suzukii. J. Insect Physiol. 99, 86–94 (2017).

37. Fisher, M. L., Fowler, F. E., Denning, S. S. & Watson, D. W. Survival of the House Fly (Diptera: Muscidae) on Truvia and Other Sweeteners. J. Med. Entomol. 54, 999–1005 (2017).

38. Gordon, H. T. Minimal nutritional requirements of the German roach, Blattella germica (L.). Ann. N. Y. Acad. Sci. 77, 290–351 (1959).

39. Özalp, P. & Emre, I. The effects of carbohydrates upon the survival and reproduction of adult female Pimpla turionellae L. (Hym., Ichneumonidae). J. Appl. Entomol. FUR Angew. Entomol. 125, 177–180 (2001).

40. Baudier, K. M. et al. Erythritol, a Non-Nutritive Sugar Alcohol Sweetener and the Main Component of Truvia®, Is a Palatable Ingested Insecticide. PLoS One 9, e98949 (2014).

41. Zheng, C., Zeng, L. & Xu, Y. Effect of sweeteners on the survival and behaviour of Bactrocera dorsalis (Hendel) (Diptera: Tephritidae). Pest Manag. Sci. 72, 990–996 (2016).

42. Burgess, E. R. & King, B. H. Insecticidal Potential of Two Sugar Alcohols to Musca domestica (Diptera: Muscidae). J. Econ. Entomol. 110, 2252–2258 (2017).

43. Burgess, E. R. & Geden, C. J. Larvicidal potential of the polyol sweeteners erythritol and xylitol in two filth fly species. J. Vector Ecol. 44, 11–17 (2019).

44. Goffin, J. et al. Toxicity of erythritol, a sugar alcohol and food additive, to Drosophila suzukii (Matsumara). Acta Hortic. 1156, 843–848 (2017).

45. Sampson, B. J., Werle, C. T., Stringer, S. J. & Adamczyk, J. J. Ingestible insecticides for spotted wing Drosophila control: a polyol, Erythritol, and an insect growth regulator, Lufenuron. J. Appl. Entomol. 141, 8–18 (2017).

46. Gilkey, P. L. et al. Lethal effects of erythritol on the mosquito Aedes aegypti Linnaeus (Diptera: Culicidae). J. Appl. Entomol. 142, 873–881 (2018).

47. Barrett, M., Caponera, V., McNair, C., O’Donnell, S. & Marenda, D. R. Potential for Use of Erythritol as a Socially Transferrable Ingested Insecticide for Ants (Hymenoptera: Formicidae). J. Econ. Entomol. 113, 1382–1388 (2020).

48. Zhang, X., Chen, S., Li, Z. & Xu, Y. Effect of Sweeteners on the Survival of Solenopsis invicta (Hymenoptera: Formicidae). J. Econ. Entomol. 110, 593–597 (2017).

49. Wentz, K., Cooper, W. R., Horton, D. R., Kao, R. & Nottingham, L. B. The Artificial Sweetener, Erythritol, Has Insecticidal Properties against Pear Psylla (Hemiptera: Psyllidae). J. Econ. Entomol. 113, 2293–2299 (2020).

50. Caponera, V., Barrett, M., Marenda, D. R. & O’Donnell, S. Erythritol Ingestion Causes Concentration-Dependent Mortality in Eastern Subterranean Termites (Blattodea: Rhinotermitidae). J. Econ. Entomol. 113, 348–352 (2020).

51. Southwick, E. E. & Pimentel, D. Energy Efficiency of Honey Production by Bees. Bioscience 31, 730–732 (1981).

52. Völkl, W., Woodring, J., Fischer, M., Lorenz, M. W. & Hoffmann, K. H. Ant-aphid mutualisms: The impact of honeydew production and honeydew sugar composition on ant preferences. Oecologia 118, 483–491 (1999).

53. Dai, Z. et al. Inter-Species Comparative Analysis of Components of Soluble Sugar Concentration in Fleshy Fruits. Front. Plant Sci. 7, (2016).

